# Bakdrive: Identifying the Minimum Set of Bacterial Driver Species across Multiple Microbial Communities

**DOI:** 10.1101/2021.09.24.461746

**Authors:** Qi Wang, Michael Nute, Todd Treangen

## Abstract

Interactions among microbes within microbial communities have been shown to play crucial roles in human health. In spite of recent progress, low-level knowledge of bacteria driving microbial interactions within microbiomes remains unknown, limiting our ability to fully understand and control microbial communities. In this study, we present a novel approach for identifying driver species within microbiomes. Bakdrive infers ecological networks of given metagenomic sequencing samples and identifies minimum sets of driver species using control theory. Bakdrive has three key innovations in this space: (i) it leverages inherent information from metagenomic sequencing samples to identify driver species, (ii) it explicitly takes host-specific variation into consideration, and (iii) it does not require a known ecological network. In extensive simulated data, we demonstrate identifying driver species identified from healthy donor samples and introducing them to the disease samples, we can restore the gut microbiome in recurrent *Clostridioides difficile* infection patients to a healthy state. We also applied Bakdrive to two real datasets, rCDI and Crohn’s disease patients, uncovering driver species consistent with previous work. In summary, Bakdrive provides a novel approach for teasing apart microbial interactions. Bakdrive is open-source and available at https://gitlab.com/treangenlab/bakdrive

## Introduction

Host-associated microbes exist as part of complex and dynamic ecosystems which have a profound impact on human health (Coyte et al., 2015; Young, 2017). Disruption of microbial communities is frequently observed after perturbations, including after antibiotic and non-antibiotic drug administration (Gibbons, 2020; Maier et al., 2018; Palleja et al., 2018; Voigt et al., 2015), and in many diseases, including recurrent *Clostridioides difficile* infection (rCDI) and inflammatory bowel disease (IBD) (Hirano and Takemoto, 2019; Menon et al., 2018; Wu and Savidge, 2020). Furthermore, several previous studies have investigated the specific ecological roles of human-associated microbes in synthetic experiments with the goal of enabling the design of beneficial microbial communities(Ji and Nielsen, 2015; Venturelli et al., 2018). Therefore, the restoration of human gut microbiota to healthy states through microbiome-based therapies, such as fecal microbiota transplantation (FMT), is a promising future therapeutic modality for improving human health (Ooijevaar et al., 2019; Wong and Levy, 2019). FMT is the most successful treatment to date in restoring microbial communities and is a clinically approved therapy for rCDI(Wang et al., 2019, Hvas et al., 2019; Ooijevaar et al., 2019). During FMT, fecal material from a healthy donor is introduced into a recipient with dysbiosis with the hope that the newly transplanted microorganisms will engraft in the recipient and jump start the restoration to a healthier community state. However, the FMT procedure is a high complexity and non-targeted intervention, and there is an inherent risk that the FMT could have negative health consequences (Davido et al., 2017; Merrick et al., 2020). Furthermore, the knowledge of the FMT mechanism is still inadequate. Which species are most important for restoring dysbiosis? How do species interact with each other during the FMT and in the new host environment? Accurate identification of keystone species in such situations remains an open research question (Banerjee et al., 2018; Röttjers and Faust, 2019). However, if the mechanisms driving the underlying biology were better understood, a community could be developed that is simpler and standardized, and that could be engrafted in the same procedure without the risk associated with FMT. It would also serve a more general purpose of improving microbiome-targeting therapeutics and pushing microbiome research closer to clinical application (Kashyap et al., 2017; Lemon et al., 2012)^,10^.

Control theory is the discipline of designing strategies to control dynamic systems (Kokotović and Arcak, 2001), and an increasingly popular application of this is in biology in the hope of manipulating or harnessing the power of microbial ecosystems (Angaroni et al., 2020; Kanhaiya et al., 2017; Thomas et al., 2019; Wolkenhauer and Mesarovic, 2005). Specifically, the members of the community are modeled as a network based on known interactions between them in the form of growth promotion or inhibition. A system is considered as controllable if it can be shifted from any initial state to a target state by applying external signals to a set of selected nodes (Cameron and Hangos, 2001). The goal of control theory in this application, specifically, is finding a minimum dominating set (MDS) of nodes, also referred to as the minimum set of drivers, that permits de facto control of the whole community. The MDS of a network is defined as the minimum set of nodes, which all the other nodes in the network directly connect to (Molnár et al., 2013). Many studies have demonstrated that an MDS can be used to find driver genes or proteins with meaningful biological functions (Kanhaiya et al., 2017; Nacher and Akutsu, 2014, 2016).

### Previous Work

Applying controllability analysis to shift microbiome states, which are characterized by microbial abundances profiles, is a field still in its infancy (Kuntal et al., 2019). Nevertheless, three recent publications have attempted to tackle the idea (Table 1). *Gibson et al*. was one of the first attempts to explore the concept, identifying “strongly interacting species’ ‘ and observing that in certain ways these are a proxy for the full community. They suggest that a small number of these species can potentially shift metagenomic samples to desired states (Gibson et al., 2016). That work predates the adoption of more common terms, like driver species, to describe such a group of microbes. Also, their analysis is strictly theoretical based on randomly generated assumptions for the interaction network. Nevertheless, they set up a framework for testing the effect of perturbations using the *Generalized Lotka-Voltera* simulation model (Hernández-Bermejo and Fairén, 1997), which has become *de rigueur* for this purpose. More recently, *Anngulo et al*. develop an algorithm to compute a minimum set of drivers given a parameterized microbial interaction network. They show that the minimum set of driver species enables controllability in the simulation environment under the assumed interaction structure (Angulo et al., 2019). Finally, *Xiao et al*. construct a model and simulation framework specifically to represent the FMT process^27^. One particular advancement over the previous simulation model is that the simulation conditions are specific to a “patient.” Namely, there is a single, parameterized global interaction network for all patients(Bashan et al., 2016). However, for a given patient, not all organisms in the global network may be present. Thus, the interaction network for that patient’s simulation is taken as the induced subgraph of the global network. The resulting diversity of simulation conditions stresses the assumptions of the previous simulation model and introduces patient-specific variability.

**Table 1.** Existing methods for identification of driver species in microbial communities. *The definition of “driver” species varies across different studies. For *Xiao et al*. we refer to beneficial species composing probiotic cocktails as “driver” species.

|  | Identifies “driver” species * | Handles multiple metagenomic samples | Does not require a known ecological network | Infers ecological networks from taxonomic classifications | Simulates FMT process | Open source, end-to-end pipeline |
| --- | --- | --- | --- | --- | --- | --- |
| *Gibson et al. (2016)* | ✓ |  |  |  |  |  |
| *Angulo et al. (2019)* | ✓ |  |  |  |  |  |
| *Xiao et al. (2020)* | ✓ |  |  |  | ✓ |  |
| ***Bakdrive, Qi et al. (2021)*** | ✓ | ✓ | ✓ | ✓ | ✓ | ✓ |

A key challenge in all three of these works noted by the authors is the difficulty of properly estimating the global interaction network parameters (Dohlman and Shen, 2019; Lv et al., 2019). This estimation for a particular host or phenotype is typically a painstaking endeavor, with many samples required to identify the strength of interactions. Another recently released software, called MICOM, took a very different approach, using a flux balance analysis to derive an interaction network from microbial genomes. It uses a broad metabolic model of the human gut microbiome and genome-wide metabolic profiles for individual microbes to impute the pairwise promotion/inhibition effects and construct the interaction network in that way (Diener et al., 2020). The effect of this is that a credible interaction network can be constructed *de novo* for each individual patient based on their community’s metagenomic composition, which would be a substantial departure from the global ecological network inherent in previous work.

An advantage of using patient-specific interaction networks, aside from sidestepping a tedious assumption, is that the range of networks across a group of patients contains information about the variance of this feature within the population, which could be useful. For the purpose of identifying a common set of driver species for the group of samples, an algorithm that can operate with multiple networks is needed. *Nacher et al*. discusses this particular problem setup, where multiple different networks with no underlying connection to one another must be controlled by a single set of driver species ^21,22^. This set of networks is represented as a multilayer network where the layers forcibly have no connection to one another, and they go on to present a heuristic for estimating a multi-layer MDS (called an MDSM), which is an NP-hard optimization.

### Assumption-Free MDS Computation

Here we present a method called Bakdrive that combines the MICOM sample-specific interaction networks with the MDSM heuristic to find a single MDS. The result is a novel method for MDS computation for a microbial community. Specifically, the networks output by MICOM for each sample are combined into a single multi-layer network, and the multilayer MDS algorithm from *Nacher et al*. (herein, simply the MDSM algorithm) is applied to compute a single MDS for the group(Nacher et al., 2019). This two-part method contains some important innovations over previous methods. For one, this method does not depend pivotally on the assumption of a single global interaction network which, while not obviously problematic, is necessarily a major simplifying assumption for both computing the MDS and simulating outcomes. Furthermore, the interaction network is computed *de novo* for each sample and thus does not need to be provided as an assumption *a priori* (Table 1).

We test Bakdrive’s ability to identify a controlling MDS using the rCDI FMT process simulation framework developed by *Xiao et al*. By varying the number of samples contributing to the multi-layer interaction network (or in other words by varying the number of layers), we show that the ability of the MDS to control the community is directly related to the size and thus diversity of the input set. We then explore the implications of a different network for every sample by exacerbating that effect: we randomly change the interaction networks for each sample by adding and removing edges, then re-run the experiments. Even with this random noise introduced, the MDS computed in this manner shows only modestly reduced effectiveness at controlling the community and remediating the rCDI condition. Finally, we compare the predictions of this model to the changes observed in an actual FMT study on rCDI patients, and we show that the simulated changes induced by the Bakdrive MDS closely resemble actual community changes in the FMT recipients.

## Results

### Simulated Driver-Species Transplantation in rCDI Patients

In this section, we investigate the impact of the number of network layers on the driver species prediction using simulated data. We first simulated 1000 pairs of diseased and healthy metagenomic samples following the protocol in *Xiao et al*.^27^. In this protocol, all the simulated samples share the same ecological network; the only difference between samples is the species abundance. After simulation, we randomly select *N* samples from the 1000 simulated healthy samples, build an *N*-layer network and use Bakdrive to find driver species. Finally, we transplant the set of driver species to the 1000 diseased samples (excluding species already present in each one) and simulate species abundances change under the GLV model. To quantify the efficacy of driver-species transplantation (briefly, DT), we compute the *recovery degree*, μ, which represents how close the *C. difficile* abundance in the after-driver transplantation (ADT) state is to that of the healthy versus diseased states, expressed as a fraction of the difference between them (here *μ*=1 represents full recovery and *μ*=0 is none).

Figure 2a shows the recovery degree for all 1000 samples after transplantation under various conditions. The panels show results using network sizes of 1, 5, 10, 20, 50, and 100. While excluding driver species that are already present in the diseased microbiota, the actual number of transplanted species varies across different samples. Additionally, for every condition, a control was run by randomly selecting an equivalent number of species to colonize. The control results are shown in green. First, it is noteworthy that although the layer-count increases from 1 to 100, the size of the MDS goes from 2 to 9. Second, the results clearly show that the MDS calculated by Bakdrive is effective at generating recovery by this measure. When *N*=100 and MDS size is 9, substantially all simulated samples experience near-full recovery.

**Figure 1.**
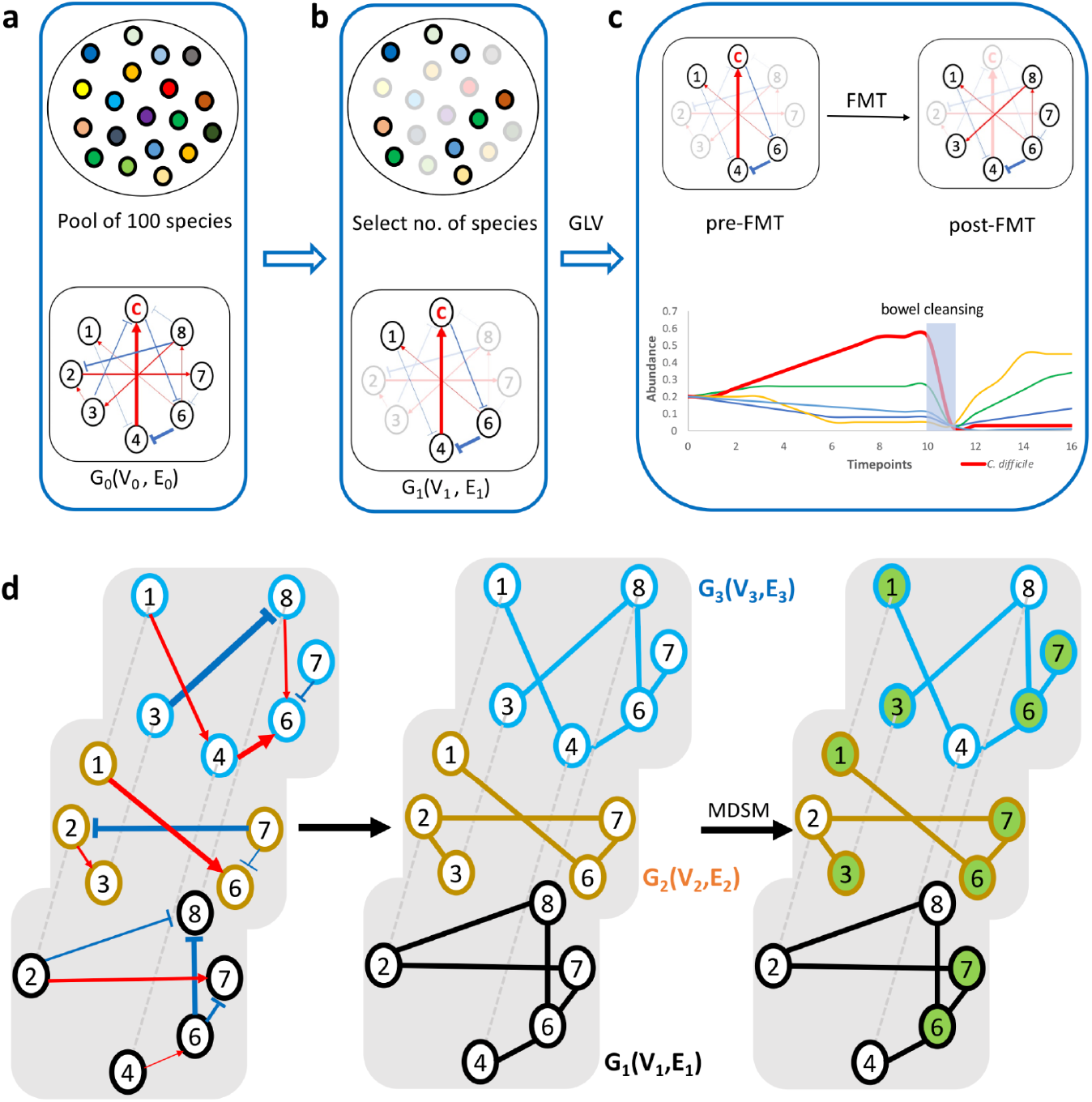
FMT process simulation and driver species identification. a The ecological network G0 of a metacommunity with 100 species. Red/blue arrows represent positive/negative relationships respectively. The widths of arrows reflect interaction strengths. b A subset of species and their corresponding interaction networks G1 are randomly selected from the pool of 100 species. c Simulation of species abundance over time. The thick red line represents the abundance of *C. difficile*. The FMT process simulation includes the pre-FMT bowel cleansing process, which leads to sudden drops of species abundance in pre-FMT samples. d Driver species identification. Directed and weighted bacteria interaction networks are converted to an undirected, unweighted multilayer network. The multilayer network consists of three layers: *G*_1_(*V*_1_, *E*_1_), *G*_2_(*V*_2_, *E*_2_), *G*_3_(*V*_3_, *E*_3_). Green nodes represent driver nodes found using the MDSM algorithm. All non-driver nodes directly connect to at least one driver node.

**Figure 2.**
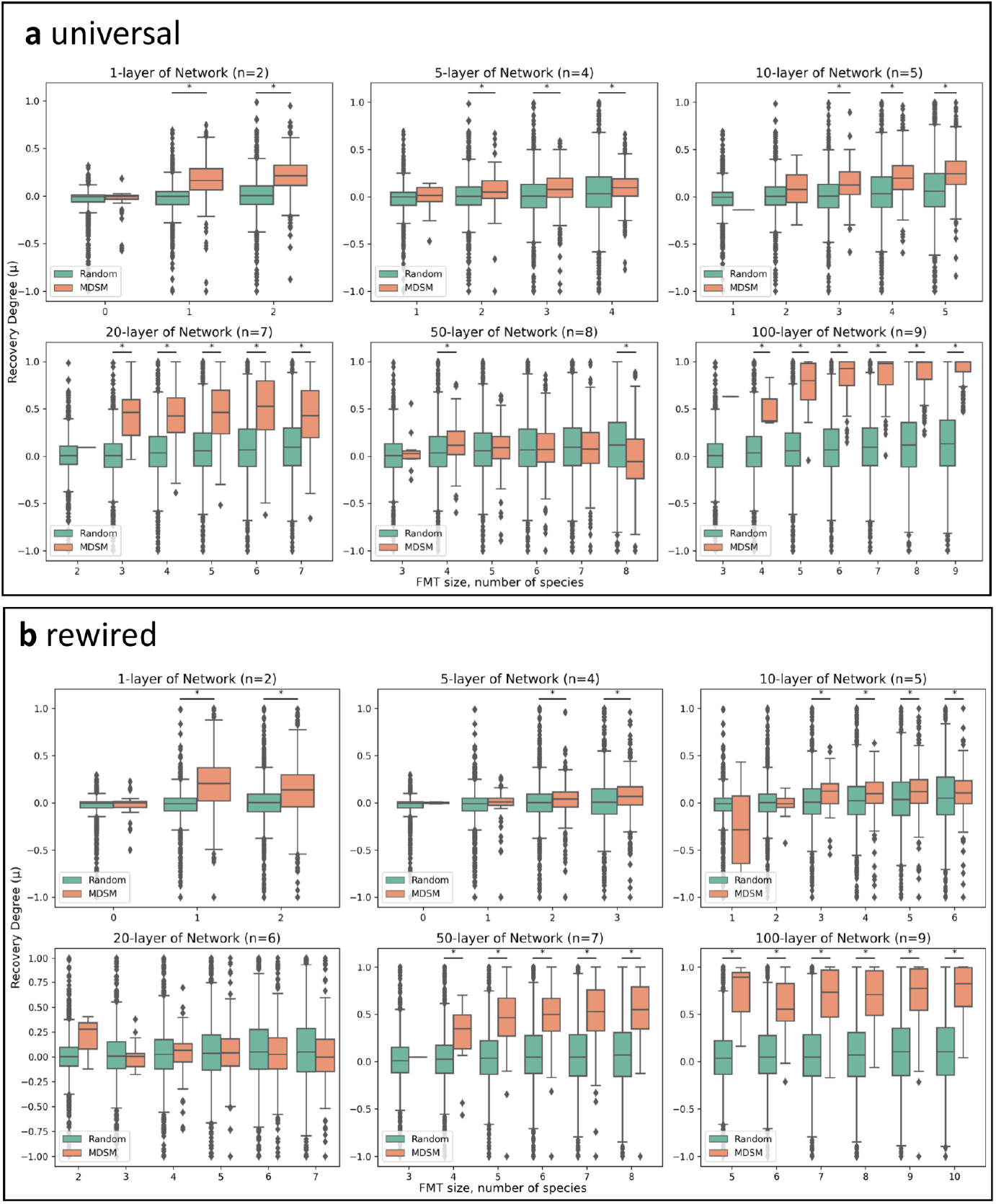
Efficacy of driver species transplantation with universal ecological networks. a) Universal b) Rewired ecological networks. We simulate the driver species transplantation process with universal/rewired microbial dynamics. In each box plot, the Y-axis shows the recovery degree of *C. difficile* after the driver species transplantation towards the 1000 simulated diseased samples. The X-axis represents the actual number of species introduced to the diseased samples. n is the number of driver species identified from a given multilayer network. The orange boxes represent the recovery degree of *C. difficile* after colonizing driver species. The green boxes represent the recovery degree of *C. difficile* after introducing the same number of species, which are randomly selected from the global pool of 100 species. The significant differences of recovery degree by Wilcoxon rank-sum test are labelled by asterisks (*, p<0.05).

Figure 3 shows this phenomenon in a slightly different manner. Here again the panels represent *N*-layer networks. Instead, we have Bray-Curtis PCoA plots of the pre-DT, post-DT, and donor samples in each case. As *N* increases, the red and blue clusters (pre- and post-transplantation, respectively) become gradually more separated from one another. In other words, not only are the samples showing normalized *C. difficile* abundance, but the overall effect on the community is becoming more pronounced with more samples.

**Figure 3.**
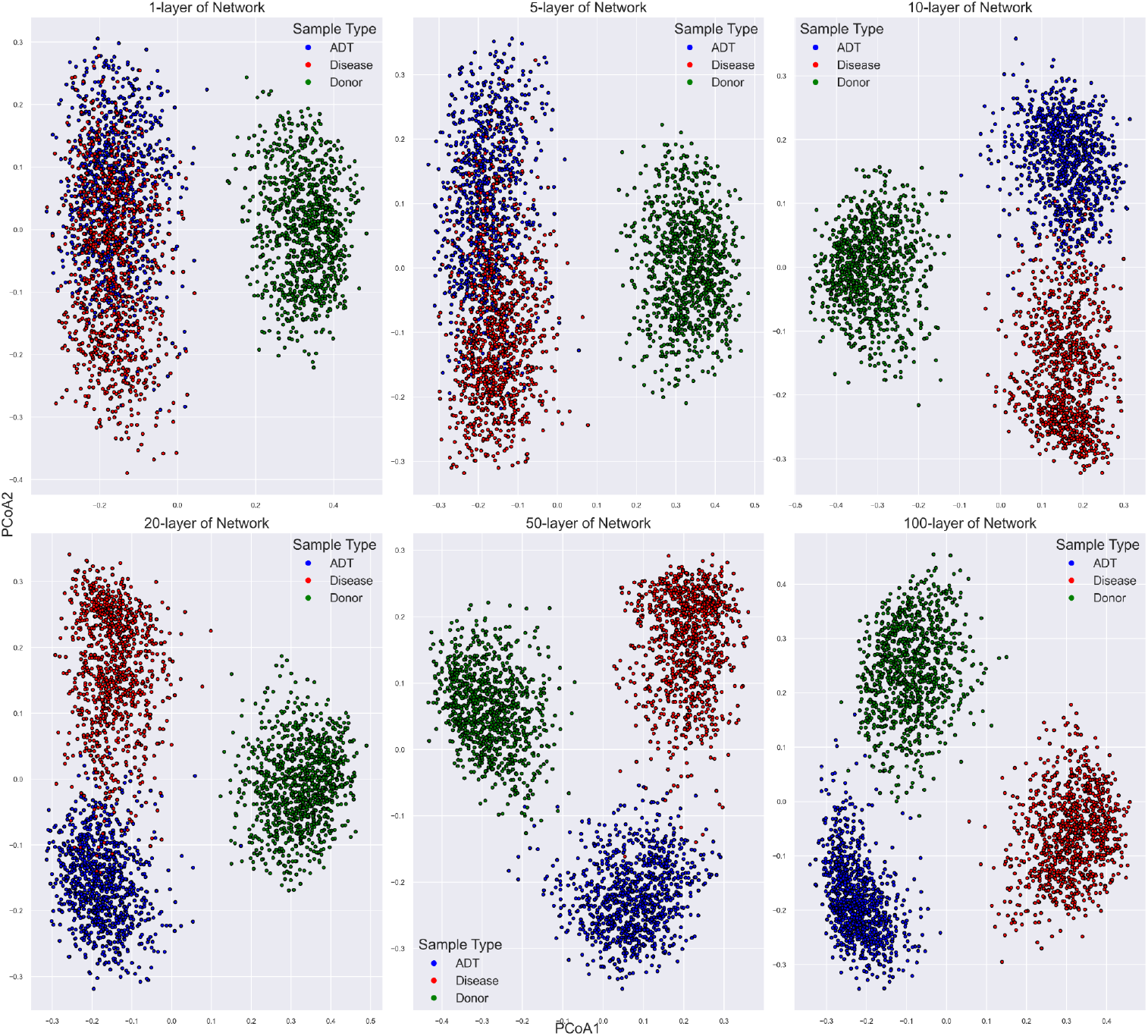
Principal Coordinate Analysis (PCoA) of simulated diseased, donor and after driver species transplant (ADT) samples. Each sample is represented as a dot. There are a total of 3000 samples: 1000 simulated diseased samples, 1000 simulated donor samples and 1000 ADT samples. Bray-Curtis dissimilarity between samples are computed.

Next, we modify the bacteria interaction networks for each simulated sample using K rewiring (Methods), which preserves the degree of each node but alters the connections between them. This random perturbation is potentially draconian but is intended to create conditions where each sample has a unique bacterial interaction network and a different intrinsic growth rate vector, an important test for Bakdrive. Results are shown in Figure 2b. Overall, the ADT efficacies of rewired bacterial interactions are consistent with the results of universal bacterial interactions, though some tempering of the recovery degree is observed. Nonetheless, using the 100-layer rewired network driver species, nearly all ADT samples were at least 50% recovered.

### Comparison with FMT-Induced Changes in Real Patients

Since the previous experiments are evaluated strictly on a simulated model of rCDI progression, and important question is, in so much as the DT leads to recovery from rCDI *in silico*, do the observed changes in microbiome composition reflect those seen in actual rCDI patients who have recovered after FMT? To evaluate this, we analyzed before, after, and donor microbiome samples from 12 rCDI patients who fully recovered after a single FMT treatment. For these patients, we apply Bakdrive using an analogous protocol: driver species are identified using the set of donor samples, then the drivers are transplanted onto the “before” community and the GLV simulations are run. The changes in community resulting from the simulated DT are then compared to what was observed in the patients during their actual recovery. This pipeline is diagrammed in additional detail in Figure 4.

**Figure4.**
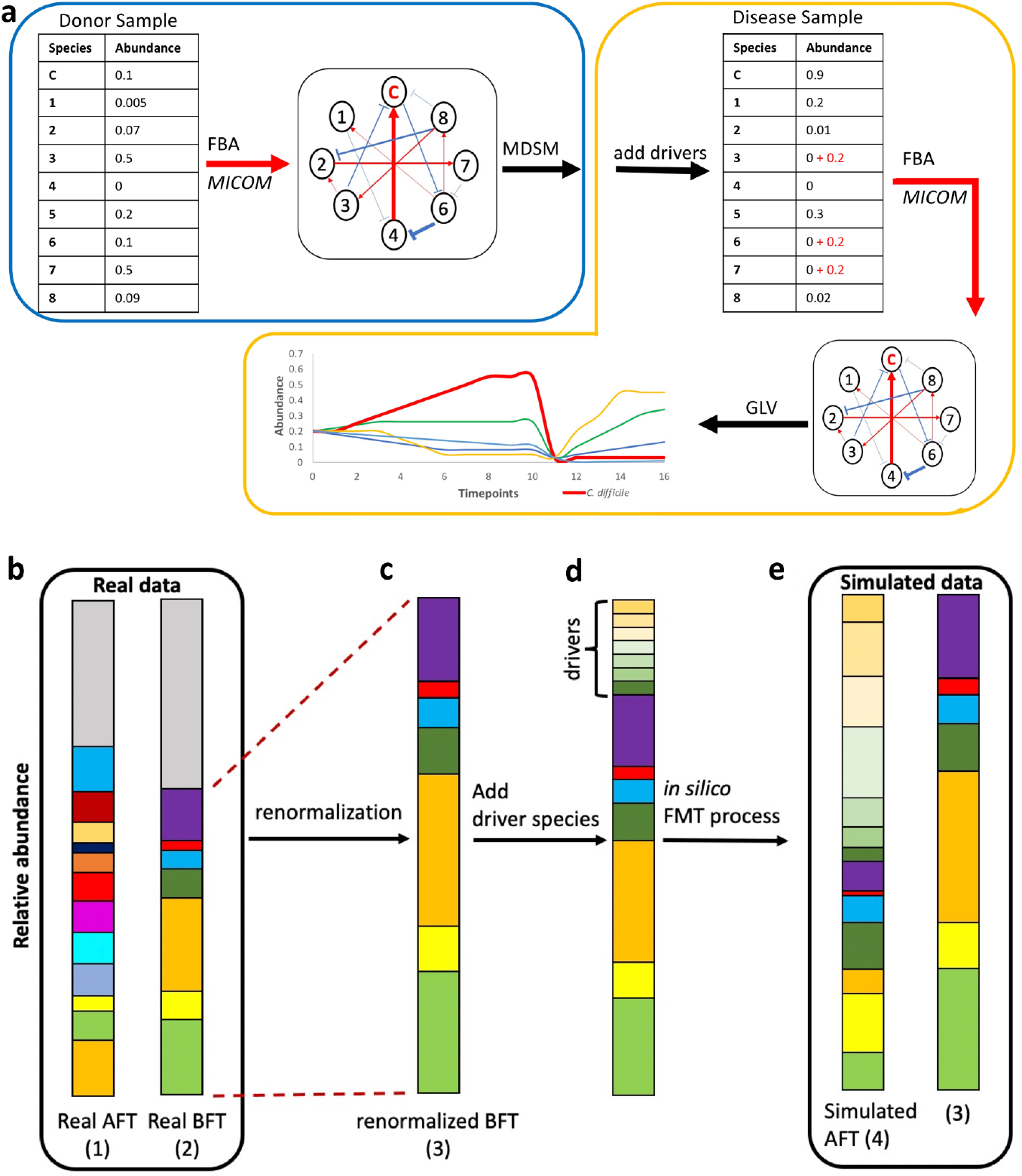
Overview of analysis on real metagenomic data. **(a)** The first step is to find driver species from a set of donor samples (blue box). The second step is to add equal amounts of driver species to patient samples, infer ecological networks of the after-FMT samples, and then simulate species abundance changes following the GLV model (yellow box). **(b-e)** Comparison between simulated and real FMT results. Each bar represents the bacterial composition of each sample. The grey area in each bar represents species without genome-scale metabolic models. b species-level taxonomic classification of real patient before-FMT (BFT) and after-FMT (AFT) samples (1,2). c relative abundances renormalization with species having metabolic models (3). d driver species and a given BFT sample combination. **(e)** FMT process simulation. Last, we compare species abundance changes between real (1,2) and simulated datasets (3,4).

We detect a total of 8 driver species from the 26 donor samples (Methods, Figure 5). Importantly, *Campylobacter jejuni* and *Streptococcus agalactiae* were identified as driver species in the donor samples and are known pathogens, and as a result were excluded from the driver-species for transplantation. This is a notable difference in the Bakdrive protocol for this experiment, but more so it is an interesting observation that may shed light on some of the risks in FMT and is discussed further in the next section.

**Figure 5.**
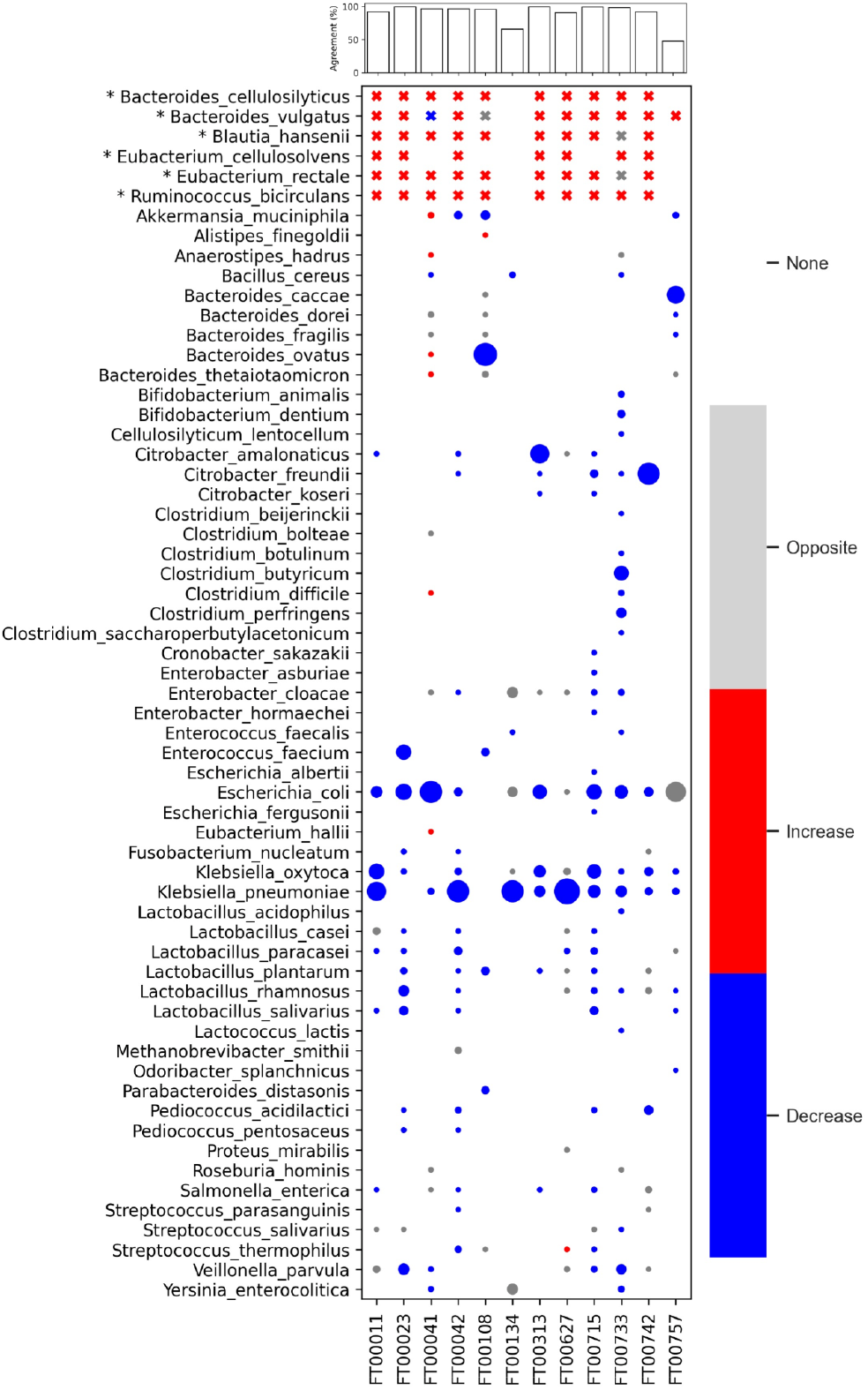
The consistency of species abundance changes between real and simulated samples. Each column represents a patient sample. Each row represents a species that exists in 12 patient samples or belongs to driver species. “*” marks driver species. The driver species abundance changes are labelled as cross, as most driver species do not exist in diseased samples. The size of the bubble represents the percentage of species in patient samples before FMT. Grey shows species abundance changes in opposite directions between simulated and real dataset. Red/blue indicates a given species abundance increases/decreases after FMT in both simulated and real data respectively. In the upper bar plot, agreement quantifies the percentage of species in renormalized diseased samples (3) whose abundances shift in the same direction in both real and simulated data.

Based on these outcomes, the question is whether the simulated changes agree with the observed changes for these patients, and the results are shown in Figure 5. Here the driver species are listed at top, followed by all other microbes in before-FMT samples. For each patient and each species, the table marker is colored based on whether abundance *increased* in both simulated and actual (red), decreased in both (blue), or changed in opposite directions (grey). The table is overwhelmingly blue and red, indicating broad agreement. For each patient, the percentage of species in agreement is also shown, and is above 90% in all but two patients, which are 45% and 65%. This is a broad indication that the simulated recoveries generated by the DT reflect the same mechanisms at work in FMT successes, a very encouraging result.

### Beyond rCDI

A major reason that most applications of control theory to the microbiome thus far have focused on rCDI is that it is arguably the only condition associated with dysbiosis where the difference between diseased and healthy community states are both simple and well-characterized. But it is most likely not the only condition where FMT, or a driver-species-based proxy for it, may turn out to be therapeutically effective. To explore the challenges this entails, we applied Bakdrive to compute sets of driver-species from a group of patients with Crohn’s disease (CD) as well as a group of healthy controls. Of interest was which species are exclusive to one set or the other. Details and results of this are given in the Supplement, section 2.1, and will be discussed further in the following section.

## Discussion

Our study focuses on the identification of driver species within and across microbial communities. Initial attempts at identifying driver species focused on inferring microbial co-occurrence patterns (Layeghifard et al., 2017; Menon et al., 2018) (Agler et al., 2016; Barberán et al., 2012; Gilbert et al., 2012); one such recent approach in this area is *Netshift (Kuntal et al., 2019)*. While these studies represent important advances specific to elucidating microbial communities, statistically inferred co-occurrences often do not reflect real interactions between bacteria (Hirano and Takemoto, 2019; Menon et al., 2018). Instead, several prior studies have suggested ecological networks calculated from biochemical reactions and metabolomics profiling can better characterize and infer bacterial interactions under different conditions (Frioux et al., 2020; Hirano and Takemoto, 2019), especially for *C. difficile (Ross et al., 2016; Wu and Savidge, 2020)*. This has led to recent focus on microbial community analysis through the lens of biochemical reactions and controllability analysis (Coyte et al., 2015). Our study is not the first to try regulating microbial communities through controllability analysis (Angulo et al., 2019; Loehle, 2006), though it differs from others in some important specifics. The major ones have been noted above: that it does not require a known interaction network, and that it uses networks that vary per sample. But an important difference from the most recent method in *Xiao et al*. is that the method presented there, in one step, uses reduction in *C. difficile* abundance as essentially a supervision criteria to select the driver-species. That makes it necessarily specific to rCDI, whereas Bakdrive is phenotype-agnostic and could possibly be applicable to additional clinical indications.

Our short sojourn into driver-species for Crohn’s disease provided some insight into Bakdrive’s broader applicability. However, beyond some similarities to other studies’ findings, evaluating whether Bakdrive has added value is not possible. Future work will focus on follow up analyses, which speculate the effect of DT on a Crohn’s patient. If this generated a positive effect on host health, an eventual outcome might be a candidate probiotic cocktail for use in a clinical trial. Theoretically, a DT is much less challenging than an FMT trial, due to the aforementioned inherent health risk of FMT therapy. However, until the disease state can be fully assessed from the community composition, *in silico* studies on other metagenome-related diseases will need to follow a different strategy than the rCDI model used in this study.

It has long been known that FMT procedures carry health risks for the recipient, a major category of which is systemic infection from pathogens acquired from the donor. An outbreak of infection from *E. coli* in March 2020 led to a change in screening procedures and an FDA advisory ^49^. In that context, the observation that driver species identified from the 12 FMT donor samples included two out of eight that were known pathogens is both alarming and, perhaps, could have been expected. In fact, the one sample with low agreement was due primarily to a sharp increase in actual abundance of *E. coli* following the FMT. That the driver species include *other* known pathogens reaffirms that FMT, for all it’s success, is far from a panacea. It also reaffirms the potential value in using Bakdrive or another contribution from control theory to identify a simpler set of microbes, which can be monitored for safety and introduced to the recipients with the same beneficial effect.

### Limitations and Future Directions

One important observation from this study is that there is no *one* optimal set of driver species for each phenotype. Driver species identification through this algorithm is influenced by many factors, including selection of input samples, accuracy of taxonomic classification, availability of species-level, genome-wide metabolic models, et cetera. For the sake of testing this, the simulation experiment described above was repeated twice with different random seeds, and even at *N*=100 layers, the set of driver species identified differs non-trivially between replicates (Table S2). That does not imply that the results themselves are unreliable, but rather that the identities of the particular species may not be as important as properties of the collective relationships to the rest of the community. But future work will ideally build on our understanding of how two largely non-overlapping sets of driver species can operate equally effectively.

Also, there are many ways in which Bakdrive, by pairing two methods designed for other contexts and applications (MICOM and MDSM), has features which are suboptimal for this particular application and considerable improvement may be possible. For example, Bakdrive does not take the directionality and strength of ecological networks into consideration. Also, the multilayer MDS problem is NP-hard^22^, so the MDSM algorithm itself is necessarily a heuristic and likely has potential for improvement in performance by itself.

Finally, a clear future direction is additional work on identifying which sets of driver species can be combined in a form of probiotic cocktail for dosing to a patient. Similar to *Xiao et al*. workflow, it suggests that the probiotic cocktail should exclude species already present in the recipient’s microbiome and there may not be one perfect set of microbes that works for all patients. It may indicate that the probiotic cocktail selection and formulation is a more involved process.

## Conclusion

In summary, Bakdrive is the first ready-to-use pipeline that can directly analyze real metagenomic sequencing data using controllability analysis and output a set of driver species that are likely to influence the direction of the entire community. Our study is the first to characterize different metagenomic states using multilayer networks and the first to do so with no *a priori* parameterization of the interaction network. The results presented here give reason to believe that the predictions from simulation might well hold up in practice, and the step of initial testing in animal models is likely close at hand. Building upon prior work, Bakdrive provides a path forward for future controllability analysis of metagenomic data and catalyzes the development of target metagenomic-based treatment.

## Methods

### Overview of Bakdrive pipeline

We implemented the Bakdrive pipeline. It has three main modules: bacterial interactions inference (interaction module), driver species identification (driver module) and FMT/DT simulation. For FMT/DT simulation, it contains three different simulations. FMT_donor module simulates the FMT process: adding donor samples directly to diseased samples, inferring bacteria interactions and simulating the after-FMT species abundance changes following the GLV model. FMT_driver module simulates the DT process: adding equal amounts of driver species to the diseased samples, inferring bacteria interactions of after-DT samples and simulating the after-DT species abundances changes. FMT_only takes the inferred bacteria interactions of after-FMT/DT samples and simulates the abundances changes directly. The pipeline is available at https://gitlab.com/treangenlab/bakdrive

### Simulation of rCDI FMT with universal dynamics

We follow the modelling procedures from *Yandong Xiao etc* paper to simulate 1000 pairs of healthy (donor) and pre-FMT diseased metagenomic samples (Xiao et al., 2020). Each simulated metagenomic sample is a random sampling from a metacommunity of 100 species. The initial abundance of sampled bacteria is 0.2. The time-dependent bacteria abundances change following the Generalized Lotka-Volterra (GLV) model with an interaction matrix A and an intrinsic growth rate vector **r** as inputs. The simulation process is built on the assumption that gut microbiomes have strong universal dynamics for healthy adults (Bashan et al., 2016; Xiao et al., 2020). Thus, all initial simulated microbial communities have the same bacteria interaction network **A** and initial growth rate **r** but with different species compositions. The interaction network **A** is also represented as a central graph, denoted as *G*_0_ (*V*_0_, *E*_0_), where each node is a species,*v*_*j*_ ∈ *V*_0_, and each edge is a negative or positive interaction between two species, {*v* _*i*_, *v*_*j*_} ∈ *E*_0 ._

After simulation, we randomly select N (1, 5, 10, 20, 50, 100) number of samples from the 1000 donor samples to build a N-layer network, and apply the MDSM approach to find driver nodes from the N-layer network (Figure1 d). After that, we introduced the driver species to the 1000 simulated diseased samples, while excluding driver species that have already existed in the diseased samples. This colonization approach has been demonstrated as the most effective FMT method in *Xiao et al*.’s study. To quantify the efficacy of DT, we borrow the concept of recovery degree, 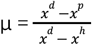, where *x*^*h*^, *x*^*d*^ and *x*^*p*^ are the abundances of *C. difficile* in healthy, diseased and after-DT states, from Xiao et al. study. We repeat each experiment of the N-layer network three times.

### Simulation of rCDI FMT with host-dependent dynamics

To test the robustness of driver nodes in suppressing pathogens, we alternate interaction networks of each simulated metagenomic sample while reserving the degree distribution of individual nodes. This practice is commonly used in complex network study. We randomly select two edges {*v* _*i*_, *v* _*j*_} and {*v* _*h*_, *v* _*k*_} from the central graph *G*_0_ (*V*_0_, *E*_0_). Meanwhile, {*v* _*i*_, *v* _*k*_} ∉ *E*_0_ and {*v* _*h*_, *v* _*j*_} ∉ *E*_0_. We add the interaction strengths and directions of{*v* _*i*_, *v* _*j*_} and {*v* _*h*_, *v* _*k*_} to {*v* _*i*_, *v*_*k*_} and {*v*_*h*_, *v*_*j*_} respectively, then delete {*v* _*i*_, *v*_*j*_} and {*v*_*h*_, *v*_*k*_} from the graph (FigureS2). We repeat this step 16 times. In this way, the degree of each node remains the same.

### MDSM algorithm

Before applying the MDSM algorithm, we need to convert a set of weighted, directed bacteria interaction networks into a multilayer network, where each layer is an undirected and unweighted bacterial interaction network *G*_*k*_ (*V*_*k*_, *E*_*k*_) of a metagenomic sample k.

The goal of the MDSM algorithm is to find a minimum set of driver species that are crucial in controlling a given metagenomic state. The metagenomic state is characterized by the multilayer network. The problem of finding driver nodes from a multilayer network is simplified as a binary integer linear programming (ILP) problem (Nacher and Akutsu, 2012).

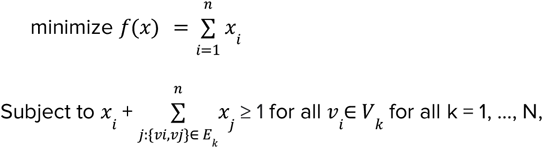

where f(x) is the dominating set and *x*_*i*_ is a binary variable of each node. *x*_*j*_ = 1 while node I belongs to the dominating set f(x). n is the total number of nodes and N is the total number of layers of a given multilayer network. Driver species identification is achieved by Bakdrive *driver* module.

### Real rCDI and CD metagenomic data analysis

The rCDI metagenomic data includes 26 donor samples from 7 donors, 19 patient samples and their corresponding after-FMT samples. Among 19 patients, 12 of them have fully recovered after receiving a single dose of FMT, while the remaining 7 patients need to receive a 2nd dose of FMT. For the simplicity of this study, we focus on analyzing the FMT results of the 12 patients with single successful FMT. The raw sequencing of rCDI is available at BioProject PRJNA454892. The sequences are classified at species level using Kraken with the full database (Wood and Salzberg, 2014).For CD, the taxonomic classification results of the HMP2 whole genome sequencing pilot dataset were directly downloaded from the IBDMDB database https://ibdmdb.org/tunnel/public/summary.html (Franzosa et al., 2018; Lloyd-Price et al., 2019). In this study, species with relative abundances below 0.1% are removed.

To identify driver species from real metagenomic data, we first construct bacteria interaction networks of individual metagenomic samples. The newly developed software MICOM provides a function of inferring bacteria interactions and species growth rate through flux balance analysis (FBA) (Diener et al., 2020). Before FBA, MICOM matches species in each sample with their genome-scale metabolic models by name. In this work, AGORA_1_03_With_Mucins is used as the model database, which contains 818 reconstructed models (Magnúsdóttir et al., 2017). If a species has multiple strains’ metabolic models in the database, we will randomly pick one as the representative model of the species. While conducting FBA, western diet is used as the default medium for growth simulation.

To infer bacteria interactions, MICOM conducts *in silico* knockout experiments and computes relative growth rate interactions between two species, denoted as *r*_*ij*_. By knocking out species i, the relative growth rate interaction of species i on j (*r*_*ij*_) is calculated. *r*_*ij*_ is defined as:

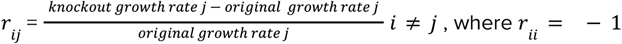

To obtain an interaction matrix for the following FMT process simulation, we need to convert *r*_*ij*_ into the interaction strength of species i on j. The meaning of interaction strengths is not constant among ecological literatures (Laska and Timothy Wootton, 1998). In this paper, interaction strength, denoted as *a*_*ij*_, is defined as competition coefficient, which measures interspecific competition relative to intraspecific competition, 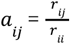. By this definition, the interaction strength *a*_*ij*_ is negative (positive) while species *i* have a negative (positive) impact on species j respectively. This step is conducted by Bakdrive *interaction* module.

### Driver species transplantation simulation of real rCDI data

Starting from estimating the absolute biomass of each species in each sample, we multiply the relative abundances of each species with estimated total human gut microbiota biomass. On average, the overall weight of human gut microbiota is 200g (Sender et al., 2016). To simulate driver species colonization of real data, we add equal amounts of driver species (default 40g/species) to the clinical patient samples (Figure 4d). After that, MICOM infers bacteria interactions and intrinsic bacteria growth rates of the after driver species transplantation (ADT) samples. It is worth mentioning that, as MICOM recommends, we divide predicted growth rates with estimated biomass in each sample, in this case, 200g of bacteria. In terms of bacterial interactions, it is hard to predict interaction strength with exact 0 value using MICOM. Thus, we assume that low interaction strengths play little role in metagenomic dynamics. If the absolute value of an interaction strength is below a given threshold, the interaction strength is set to 0, that is, we assume there is no bacterial interaction between the two species. To choose a reasonable interaction strength threshold, we create a screen plot, where x axis is a list of absolute values of interaction strengths and y axis is corresponding 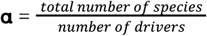. Based on empirical experiments, the threshold at the inflection point gives the best performance. In this study, we select 0.2 as the threshold for both C. difficile and IBD datasets (FigS3). After adjusting growth rates and bacteria interaction strengths, we simulate the population dynamics following the GLV-based modeling framework proposed by *Xiao et al*. study (Xiao et al., 2020). The simulation stops at steady state or the last time point all the species have positive abundances. This step is performed by Bakdrive *fmt_driver* module.

### Simulated and real rCDI FMT process comparison

To quantify the efficacies of simulated driver species transplantation, we calculate the percentage species in renormalized BFT samples whose abundances move in the same directions in simulated and real after FMT process. We also refer to this value as agreement. To be clear, we only examine species with metabolic models in AGORA database, that is, whose bacteria interactions can be estimated by MICOM. We first compute species abundance changes between patients’ BFT samples and their AFT samples from the real data (Magnúsdóttir et al., 2017; Park et al., 2019). To compare the abundance shfits between real and simulated data, we renormalize the relative abundance of species in BFT samples with species who have metabolic models in AGORA database. We then calculate species abundance changes between re-normalized BFT samples and simulated ADT samples (Figure 4).

In terms of agreement calculation, if the relative abundance of species i shifts in the same direction in both simulated and real BFT and AFT samples of a given patient, we consider species i as having consistent abundance movement. We then added the percentage of species i in renormalized BFT to the agreement of the patient.

### Statistical Analysis

Bray-Curtis similarities and principal coordinate analysis (PCoA) is conducted using skbio.diversity and skbio.stats.ordination packages(The scikit-bio development team, 2020). Statistical significance testing is decided by Wilcoxon rank-sum using the scipy.stats.ranksums package(Virtanen et al., 2020). The script is available at: https://gitlab.com/treangenlab/bakdrive/-/tree/data

## Acknowledgements

Q.W. was supported by funds from Rice University and by funds from the National Institute for Neurological Disorders and Stroke (NINDS) of the National Institutes of Health under award number R21NS106640. M.N. was funded by a fellowship from the National Library of Medicine Training Program in Biomedical Informatics and Data Science (T15LM007093, PI: Kavraki). T.J.T was partially supported by funds from NIH award P01-AI152999..

## Author Contributions

Q.W. and T.J.T. conceived and managed the study. All authors (Q.W., M.N., T.J.T.) designed the experiment(s) and analyzed the results. Q.W. conducted the experiment(s). Q.W., M.N., and T.J.T. wrote the manuscript. All authors (Q.W., M.N., T.J.T.) reviewed and approved the final manuscript.

## Competing interests

The authors declare no competing interests.

## Supplementary Notes

### 1. Supplementary Methods

#### 1.1 MICOM

MICOM is a software designed to predict growth rates of bacterial species inside the gut and infer metabolic interactions within the microbial communities. MICOM first takes genus- or species-level classifications for each metagenomic sample and matches organisms with the AGORA genome-scale metabolic models by name. MICOM then conducts flux balance analysis (FBA) with L2 regularization. FBA approximates steady-state metabolic fluxes of a given organism from its genome-scale metabolic model by maximizing the production of organism-specific biomass under constraints. For metagenome analysis, FBA is often used to maximize community growth rate. However, by simply maximizing the community growth rate, it ignores individual growth rate optimization and removes many existing taxa from the microbial community. Therefore, to make sure that bacteria present in a metagenomic sample can continually grow, MICOM performs FBA coupled with L2 regularization. L2 regularization assists FBA to optimize individual and community growth rates simultaneously, so that more organisms are able to grow. Also, MICOM prediction takes dietary and taxon abundances into consideration, in order to generate personalized metabolic interactions for individual metagenomic samples.

Additionally, MICOM can be used to simulate a species knockout by removing the species from the following FBA. The bacteria interactions are inferred from the growth rate before and after the knockout. Given species j knocked out from the environment, if the growth rate of species i increases (decreases), species j have a negative (positive) impact on species i, respectively. The relative growth rate changes before and after knockout can be used to calculate interaction strength between the two species.

#### 1.2 rCDI metagenome simulation using the GLV model

Our rCDI metagenome simulation follows the protocol *Yandong Xiao et. al*. proposed. *Yandong Xiao et. al*. conduct the FMT simulations by taking N species following the Generalized lotka-Volterra (GLV) model with an interaction matrix A and an intrinsic growth rate vector r. First, we generate the interaction matrix A and intrinsic growth rate vector r. The interaction matrix is constructed using a directed Erdos-Rennyi random graph model with N nodes and C connectivity, that is, two species have C probability to connect with a directed edge. The interaction strength *a*_*ij*_ is drawn from a normal distribution. In addition, a negative self loop, *a*_*ij*_ = -1, is added to the system. For the intrinsic growth rate vector r, the value of *r*_*i*_ is drawn from a uniform distribution ∼U[0,1].

In order to simulate healthy gut microbiota, the FMT simulation method first selects a subset of species from the metacommunity with its species richness above a given threshold. The initial abundance profile is chosen from the uniform distribution U[0,1]. For population dynamics of simulated microbiota communities, we first follow the universal dynamics. In this case, all samples share the same interaction matrix A and an intrinsic growth rate vector r. The only difference between healthy metagenomic samples is the set of species present in the communities. For non-universal dynamics, each community has its own interaction matrix A and an intrinsic growth rate vector r. The interaction matrix is derived from the initial interaction matrix *A*_0_ and rewired following the degree-preserving rewriting method.

Given the interaction matrix A and an intrinsic growth rate vector r for each sample, we simulate their population dynamics following the GLV model until the system reaches a steady state. If the steady state abundance of *C. difficile* is less than 10^−5^and the number of species is above 60 (total 100 species), we keep it as a healthy community. Otherwise, we discard it.

On the other hand, a diseased metagenomic sample is derived from a healthy local community by randomly removing part of its species, which is designed to mimic the impact of antibiotics administration. We then simulate the population dynamics until it reaches a steady state. As long as the abundance of *C. difficile* is larger than default threshold 0.5, the sample is considered as diseased.

For driver species transplantation simulation, we combine a diseased community with driver species by directly adding the abundances of driver-specific species to the diseased samples. We then simulate the population dynamics until the newly merged system reaches a steady state.

#### 1.3 Generalized lotka-Volterra (GLV) model

The GLV model is used to recapitulate direct competitive and cooperative relationships between any number of species(Hofbauer et al., 1998). In this case, the GLV model simulates the dynamics of N bacteria inside the gut microbiome. The population is a vector x and the GLV model contains a set of ordinary differential equations given by

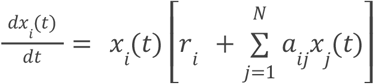

*x*_*i*_ (*t*) represents the abundance of species i at time t; *r*_*i*_ represents the intrinsic growth rate of species i. *a*_*ij*_ denotes the interaction strength between species i and j, that is, the direct impact of species j on i.

### 2. Supplementary Results

#### 2.1 Crohn’s Disease

The importance of microbiota in the intestinal inflammation of Crohn’s disease (CD) has been long recognized. In general, an observed dysbiosis in CD patients has lower abundances of *Clostridium cluster IV / XIVa* and higher abundances of *Bacteroides* (Joossens et al., 2011). According to our analysis, there are a total of 25 driver species in the CD and 23 driver species in the nonIBD samples. Among them, 15 driver species are present in both groups (Table S1). The unique driver species of CD are enriched in Bacteroides and the unique nonIBD driver species are concentrated in *Clostridium cluster IV / XIVa*. The nonIBD/CD driver species distributions at genus level are consistent with previous findings (Joossens et al., 2011). On species level, the unique CD driver species *Parasutterella excrementihominis (Chen et al., 2018; Joossens et al., 2011)* and *Veillonella parvula (Bajer et al., 2017; Gevers et al., 2014; Joossens et al., 2011)* have been shown to have distinct associations with CD or intestinal inflammation. On the other hand, the unique nonIBD driver species Bifidobacterium adolescentis and Dialister invisus are frequently observed to have decreased presences in CD metagenomic samples (Joossens et al., 2011).

Despite the success in detecting driver species associated with CD, CD signature species are not always identified as unique driver species of CD. For example, the decrease of *Faecalibacterium prausnitzii* has been considered as signals of dysbiosis of CD (Jia et al., 2010; Joossens et al., 2011; Sokol et al., 2008). However, it is not identified as unique drivers of nonIBD, but drivers of both CD and nonIBD groups. Without considering the oversights of the pipeline, the discrepancies between abundance changes analysis and driver species identification indicate that controllability analysis can potentially provide a new insight in understanding the importance of bacteria in different metagenomic states.

#### 2.2 Supplementary Figures and Tables

**Figure S2.**
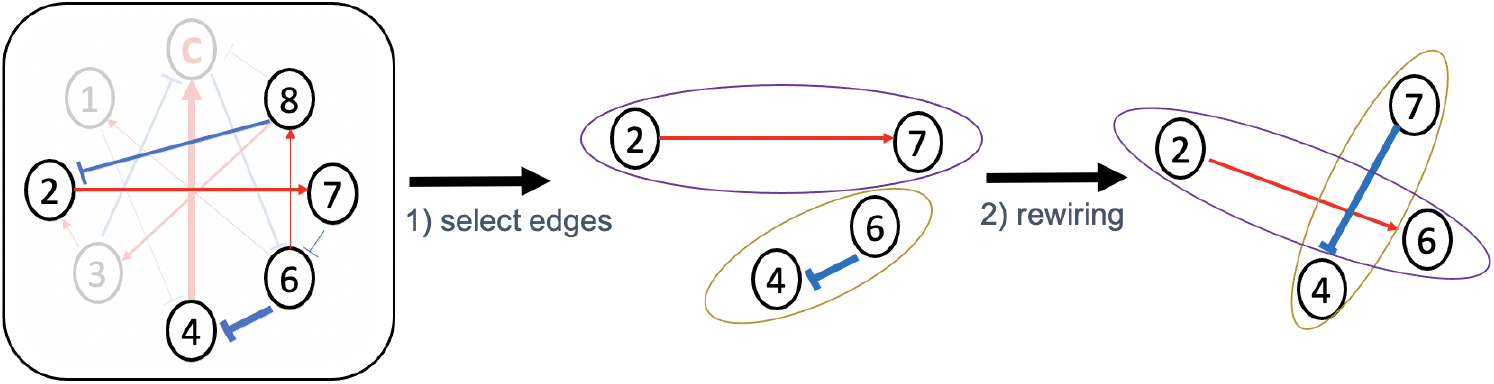
Degree-preserving rewiring method. We randomly select two edges, in this case {2, 7} and {6, 4}, from the central graph*G*_0_. We transfer the edge values of {2, 7} and {6, 4} to {2, 6} and {7, 4} respectively.

**Figure S3.**
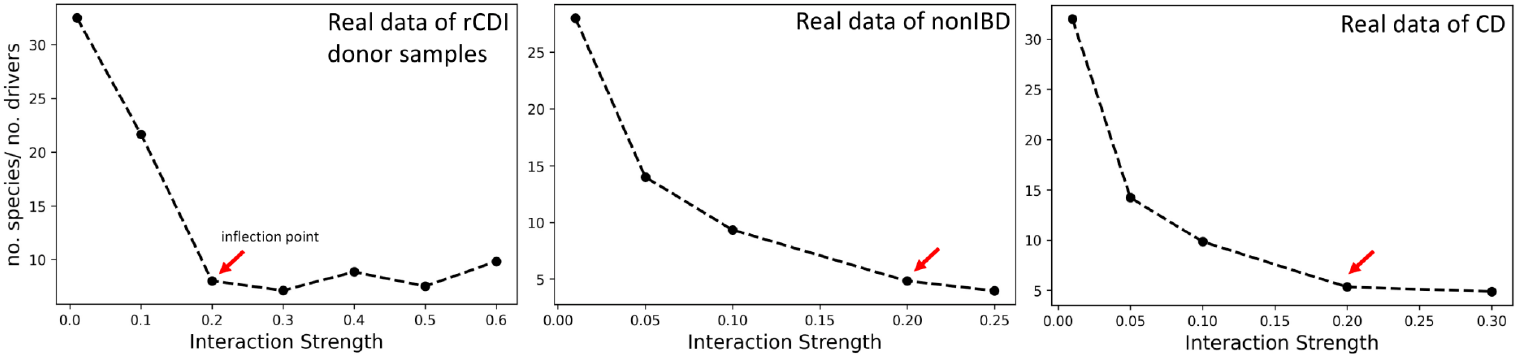
Interaction strength threshold selection. X-axis is the absolute value of interaction strength threshold. The edges with the interaction strength below the threshold are removed from ecological graphs. Y-axis represents 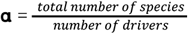. Based on empirical experiments, the interaction strength threshold at the inflection point gives the best performance.

**Table S1.**
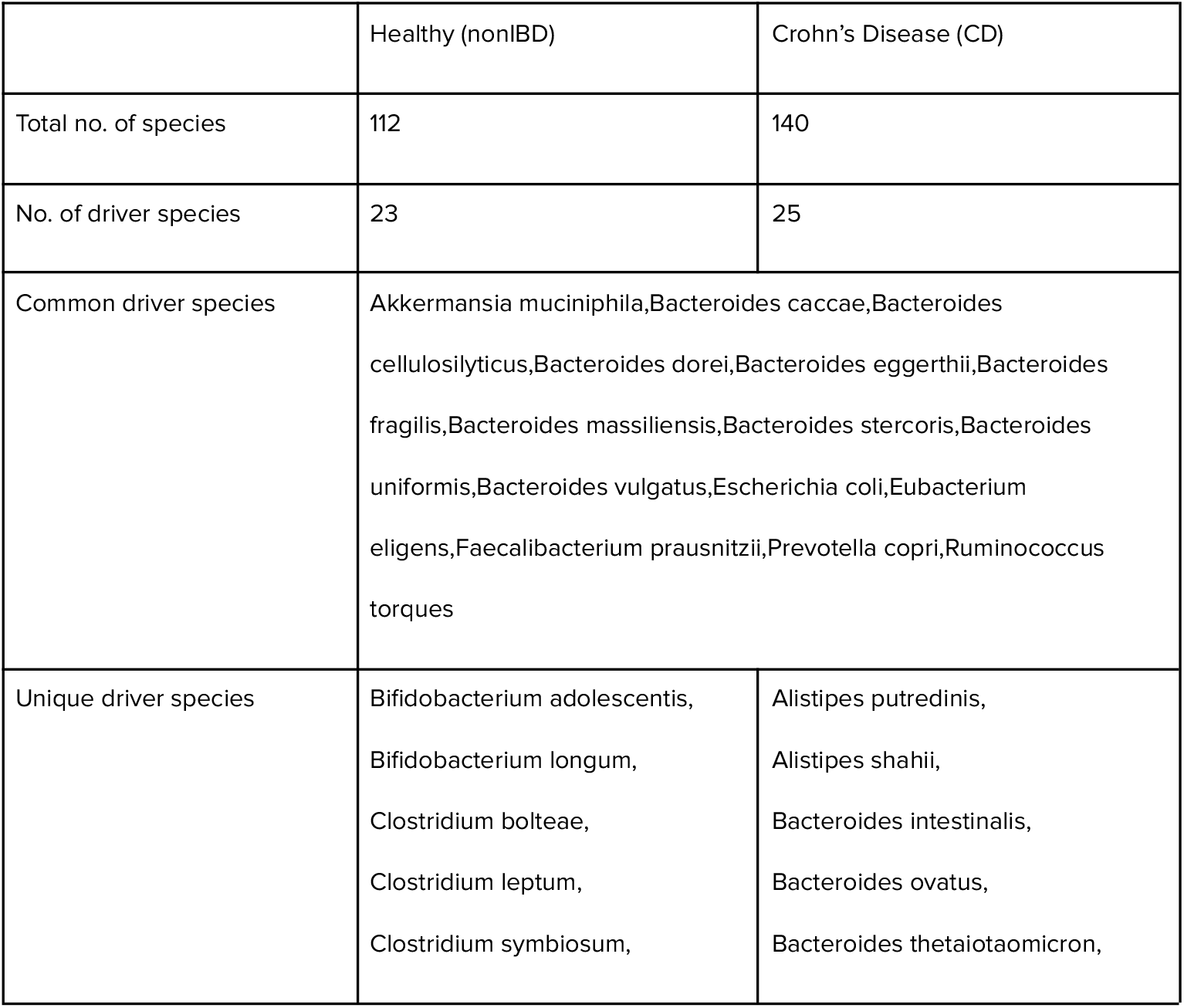
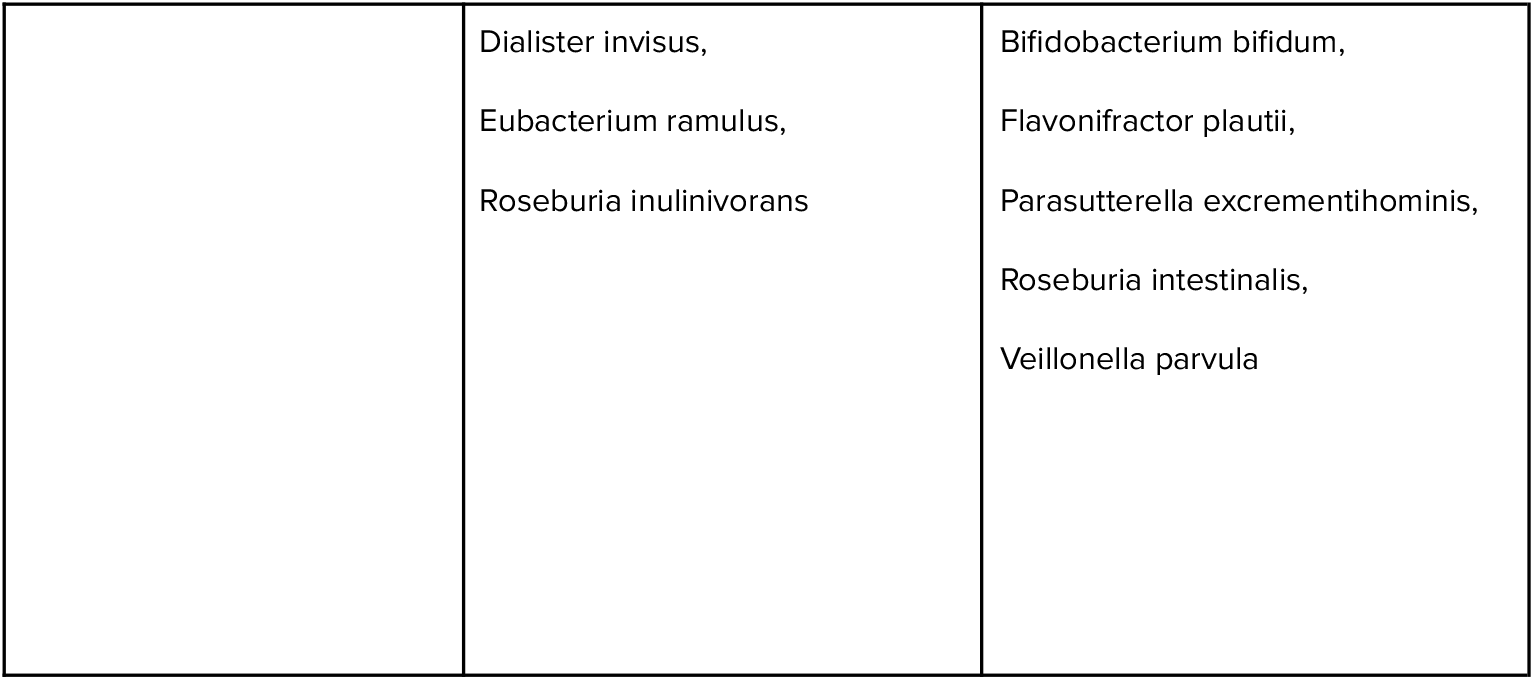
Driver species of nonIBD and CD metagenomic samples.

**Table S2.** Driver nodes of simulated data with universal ecological networks.

| No. of layers | No. of drivers | Driver Nodes Id |  |  |
| --- | --- | --- | --- | --- |
|  |  | Repeat1 | Repeat2 | Repeat3 |
| 1 | 2 | 56,92 | 60,68 | 86,92 |
| 5 | 4 | 4,33,62,63 | 60,64,68,79 | 52,53,63,76 |
| 10 | 5 | 22,61,71,73,82 | 26,36,38,60,68 | 7,53,76,92,99 |
| 20 | 5-7 | 14,25,27,60,68,87,94 | 22,26,54,93,94 | 53,58,76,82,93,99 |
| 50 | 8-9 | 27,46,51,50,60,62,68,73,8<br>2 | 3,30,46,54,62,63,9<br>2,94 | 7,14,15,22,27,50,61,92 |
| 100 | 9 | 14,22,25,3,30,54,80,92,94 | 7,22,33,46,60,69,9<br>2,93,94 | 14,22,25,27,46,54,64,8<br>4,92 |

## Notes

https://gitlab.com/treangenlab/bakdrive

### Competing Interest Statement

The authors have declared no competing interest.

